# Multiple large-scale gene and genome duplications during the evolution of hexapods

**DOI:** 10.1101/253609

**Authors:** Zheng Li, George P. Tiley, Sally R. Galuska, Chris R. Reardon, Thomas I. Kidder, Rebecca J. Rundell, Michael S. Barker

## Abstract

Polyploidy or whole genome duplication (WGD) is a major contributor to genome evolution and diversity. Although polyploidy is recognized as an important component of plant evolution, it is generally considered to play a relatively minor role in animal evolution. Ancient polyploidy is found in the ancestry of some animals, especially fishes, but there is little evidence for ancient WGDs in other metazoan lineages. Here we use recently published transcriptomes and genomes from more than 150 species across the insect phylogeny to investigate whether ancient WGDs occurred during the evolution of Hexapoda, the most diverse clade of animals. Using gene age distributions and phylogenomics, we found evidence for 18 ancient WGDs and six other large-scale bursts of gene duplication during insect evolution. These bursts of gene duplication occurred in the history of lineages such as the Lepidoptera, Trichoptera, and Odonata. To further corroborate the nature of these duplications, we evaluated the pattern of gene retention from putative WGDs observed in the gene age distributions. We found a relatively strong signal of convergent gene retention across many of the putative insect WGDs. Considering the phylogenetic breadth and depth of the insect phylogeny, this observation is consistent with polyploidy as we expect dosage-balance to drive the parallel retention of genes. Together with recent research on plant evolution, our hexapod results suggest that genome duplications contributed to the evolution of two of the most diverse lineages of eukaryotes on Earth.

## Introduction

Genome duplication has long been considered a major force of genome evolution and a generator of diversity. Evidence of paleopolyploidy is found in the genomes of many eukaryotes, such as yeasts, teleost fishes, and plants (1–4). Polyploid speciation is perhaps most important among plants where nearly ⅓ of contemporary vascular plant species have recently duplicated genomes (5, 6). All extant seed plants have also experienced at least one ancient WGD (7, 8), and many flowering plants have undergone multiple rounds of paleopolyploidy (2, 3, 9). The creation of new genes (10, 11), higher turnover of genome content (12, 13), and increased rates of adaptation (14) following polyploidy have likely contributed to the diversification of flowering plants (12, 15).

In contrast to plants, polyploid speciation among animals is generally regarded as exceptional (16, 17). The most well known polyploidization events in animals are two rounds of ancient WGD (the 2R hypothesis) that occurred in the ancestry of all vertebrates (18, 19). However, most known cases of polyploidy in animals are found among parthenogenetic and hermaphroditic groups (17, 20). If paleopolyploidy is indeed fundamental to the evolution of animal life across deep time, as it is in plants, we would expect to find WGDs throughout the most species-rich animal lineages: molluscs and arthropods. Little is known about ancient WGD among invertebrates, but there is growing evidence for paleopolyploidy in molluscs (21) and chelicerates (22–24). There is no evidence of paleopolyploidy among Hexapoda, the most diverse lineage of animals on Earth. Only 0.01% of the more than 800,000 described hexapod species (25) are known polyploids (17, 20). However, until recently there were limited data available to search for evidence of paleopolyploidy among the hexapods and other animal clades. Thus, the contributions of polyploidy to animal evolution and the differences with plant evolution have remained unclear.

To search for evidence of WGDs among the hexapods, we leveraged recently released genomic data for the insects (26). Combined with additional data sets from public databases, we assembled 128 transcriptomes and 27 genomes with at least one representative from each order of Hexapoda (SI Appendix, Dataset S1). We selected data from chelicerates, myriapods, and crustaceans as outgroups. Ancient WGDs were initially identified in the distributions of gene ages (Ks plots) produced by DupPipe (27, 28). We also used the MAPS algorithm (8) to infer WGDs or other large-scale genome duplications that are shared among descendant taxa. MAPS uses multi-species gene trees to infer the phylogenetic placement of significant bursts of ancient gene duplication based on comparison to simulated gene trees with and without WGDs. Simulations were conducted with GenPhyloData (29) with background gene birth and death rates estimated from WGDgc (30) for each MAPS analysis (Dataset S3-S4). Analyses of synteny within the *Bombyx mori* genome (31) provided additional evidence that significant duplications inferred by our MAPS analyses may result from large-scale genome duplication events. We also compared the synonymous divergence of putative WGD paralogs with the orthologous divergence among lineages to place inferred genome duplications in phylogenetic context. Potential ancient WGDs detected in our gene age distributions were further corroborated by analyses of biased gene retention across 20 hexapod genomes.

## Results

### Inference of WGDs from Gene Age Distributions

Our phylogenomic analyses revealed evidence for WGDs in the ancestry of many insects. Peaks of gene duplication consistent with WGDs were observed in the gene age distributions of 20 hexapod species (Fig. 1, SI Appendix, Fig. S1-S4, and Table S1). Each of the inferred WGDs was identified as a significant peak using SiZer and mixture model analyses (SI Appendix, Fig. S1, Table S1, Dataset S2). Fifteen of these appear as phylogenetically independent WGDs because the sampled sister lineages lack evidence of the duplications (Fig. 2, SI Appendix, Fig. S1-S4). In two cases, multiple sister lineages contained evidence for paleopolyploidy in their Ks plots. All sampled species of Thysanoptera contained evidence of at least one peak consistent with paleopolyploidy in their Ks plots (Fig. 1*B*, SI Appendix, Fig. S2*I*-*K*). Analyses of orthologous divergence among these taxa indicated that the putative WGD peaks are older than the divergence of these lineages, and we currently infer a single, shared WGD in the ancestry of Thysanoptera. Similarly, multiple taxa in the Trichoptera had evidence for WGD(s) (SI Appendix, Fig. S1*O*-*R*, and Fig. S2*O*-*R*). Analyses of orthologous divergence indicated that each of these putative WGDs occurred independently (SI Appendix, Fig. S5*O* and *P*, and Table S2). A MAPS analysis also supported the independence of these WGDs and found evidence for a deeper duplication event shared among all the sampled Trichoptera (SI Appendix, Fig. S6*Y* and Table S3).

**Fig. 1.**
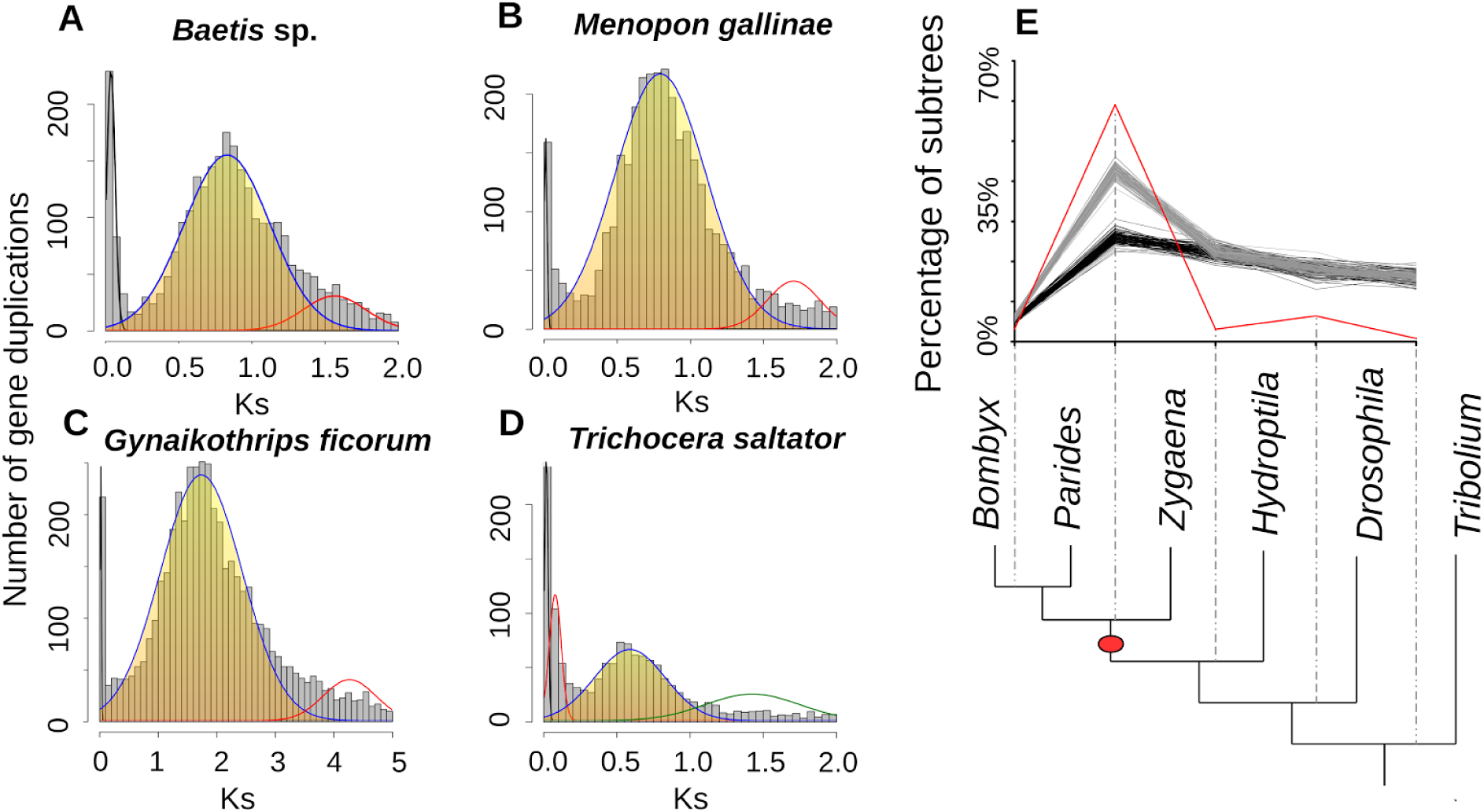
Inferring ancient WGDs and large-scale genome duplications. Histograms of the age distribution of gene duplications (Ks plots) with mixture models of inferred WGDs for (**A**) *Baetis* sp. (Ephemeroptera); inferred WGD peak median Ks = 0.83. (**B**) *Gynaikothrips ficorum* (Thysanoptera); inferred WGD peak median Ks = 1.73. (**C**) *Menopon gallinae* (Psocodea); inferred WGD peak median Ks = 0.80. (**D**) *Trichocera saltator* (Diptera); inferred WGD peak median Ks = 0.59. The mixture model distributions consistent with inferred ancient WGDs are highlighted in yellow. (**E**) MAPS results from observed data, null and positive simulations on the associated phylogeny. Percentage of subtrees that contain a gene duplication shared by descendant species at each node, results from observed data (red line), 100 resampled sets of null simulations (multiple black lines) and positive simulations (multiple gray lines). The red oval corresponds to the location of an inferred large-scale genome duplication event in Lepidoptera.

**Fig. 2.**
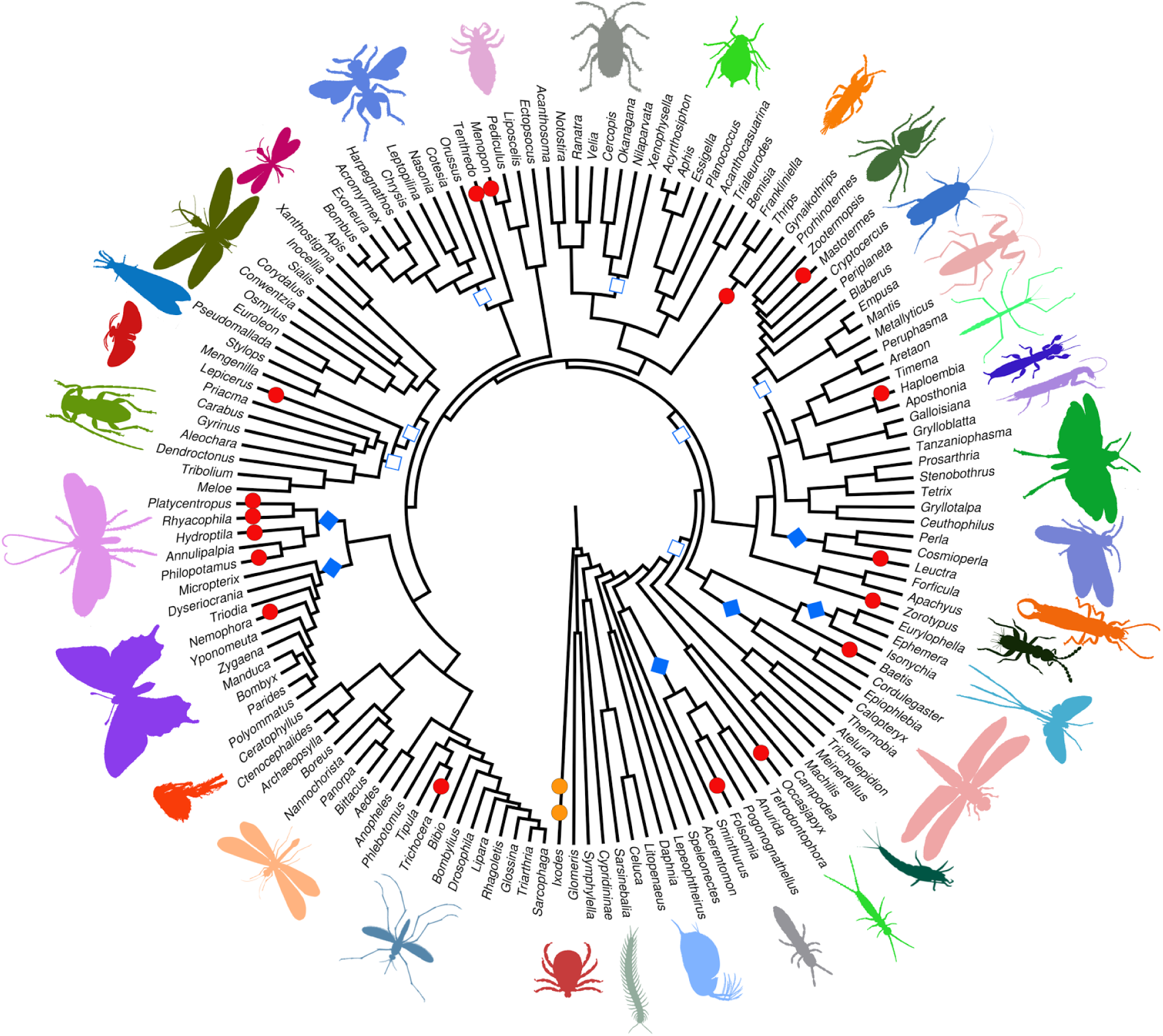
Placement of inferred ancient genome duplications on the phylogeny of Hexapoda. Red circles = WGDs in hexapods inferred from Ks plots; Orange circles = WGDs in outgroups inferred from Ks plots; Blue diamonds = large-scale genome duplications inferred by MAPS analyses; Empty squares = episodic bursts of gene duplication with varying levels of significance across different MAPS analyses. Hexapod phylogeny adapted from Misof et al. 2014. The *Solenopsis invicta* (Hymenoptera) WGD inferred by Ks plot is not included on this phylogeny. Images of Raphidioptera, Coleoptera, and Neuroptera credit to Tang Liang, Zichen Wang and Zheng Li. Other images are in the public domain (Table S6).

Overall, our analyses of gene age distributions found evidence for 18 independent paleopolyploidizations in the ancestry of 14 orders of hexapods (Fig. 2, SI Appendix, Fig. S7). We observed evidence for ancient WGDs in diverse lineages of hexapods including springtails, beetles, ants, lice, flies, thrips, moths, termites, sawflies, caddisflies, stoneflies, and mayflies. Some of these WGDs were of relatively modest synonymous divergence and may be correlated with the origins of families or clades of genera, such as the inferred paleopolyploidization in *Trichocera saltator* (Fig. 1*D*). However, many of these putative WGDs appear to have occurred early in the evolution of different hexapod orders with relatively high synonymous divergence among paralogs. For example, applying the *Drosophila* synonymous substitution rate of 5.8 × 10^−9^ substitutions/synonymous site/year (32) to thrips, we estimated that the thrips duplication occurred approximately 155 MYA based on the median paralog divergence of the WGD. However, if thrips have a slower rate of evolution than *Drosophila*, then this WGD would be older.

### Phylogenomic Inference and Simulation of Ancient Large-scale Genome Duplications

Given the depth of the phylogeny, there may be many WGDs or other large-scale genome duplications in the ancestry of hexapods that do not appear in Ks plots due to saturation of substitutions. We conducted 33 MAPS (8) analyses of 111,933 nuclear gene family phylogenies to infer large bursts of gene duplication deep in the history of all major clades of hexapods (SI Appendix, Fig. S6 and Dataset S3). By examining shared gene duplications from multiple species, MAPS increases the signal of deep duplications and provides more resolution than a single species analysis. Overall, we found 25 branches in 22 MAPS analyses that contained at least one branch with significantly more shared gene duplications than expected compared to the null simulations (SI Appendix, Fig. S8 and Dataset S3). To further characterize these significant gene bursts, we simulated an additional set of gene trees with a WGD at the phylogenetic location of the duplication bursts in these 22 analyses. We found that 14 of the 25 bursts of gene duplication were statistically consistent with our positive simulations of WGDs (SI Appendix, Fig. S9 and Dataset S4). Six of these large-scale genome duplications had robust and consistent evidence across all MAPS analyses (Fig. 2, SI Appendix, Fig. S6, Table S3 and Dataset S3-S4). These included large-scale genome duplications in the ancestry of Odonota, Lepidoptera, and Trichoptera. We also observed seven episodic bursts of gene duplication in the insect phylogeny that had varying levels of significance across different MAPS analyses (eg. Coleoptera, Hymenoptera (in part) and Hemiptera (in part); Fig. 2, SI Appendix, Fig. S6-7 and Dataset S4). Although there was conflict among our analyses as to whether these seven events were statistically consistent with large-scale genome duplications, they do reflect significant increases in gene duplication at these locations in the insect phylogeny.

Considering the putative ancient nature of the MAPS inferred duplications, most are likely too saturated to appear in any of our Ks plots. Saturation of substitutions diminishes the signature peaks of polyploidy in Ks plots, and WGDs with a peak of Ks > 2 may become difficult to detect (28, 33, 34). To confirm that we should not expect to see these six large-scale genome duplications in Ks plots, we tested if the ortholog divergence among these lineages was Ks > 2. As expected, the median ortholog divergence was Ks > 2 (SI Appendix, Fig. S5 and Table S2). Thus, all six of the large-scale genome duplications inferred with MAPS are likely too saturated to appear in Ks plots.

Synteny provides perhaps the most compelling evidence to characterize ancient genome duplication events. Although there are a number of high quality hexapod genomes, there are few high quality genomes in the clades where we inferred ancient duplications. The genome of *Bombyx mori (31)*, the silkworm moth, is one of the few genomes that is reasonably well assembled and has an ancient large-scale genome duplication based on our MAPS analyses (SI Appendix, Figs. S8*AA* and S9*AA*, Dataset S5). If this ancient duplication involved structural duplications, as with a WGD or other large-scale chromosomal duplication event, then we expect to find a significant association of syntenic chains with the MAPS paralogs in the genome of *B. mori*. Using CoGe’s SynMap tool (35), we identified 728 syntenic chains that included 2210 genes (SI Appendix, Fig. S11). To test the significance of this association, we used a statistical approach similar to a method developed to find evidence of paleopolyploidy from synteny in linkage mapping data (36). We found that significantly more syntenic chains—83 chains—were associated with our MAPS paralogs than expected by chance (chi-square test *p*-value = 0.0001). Although many of these syntenic chains are small, this is not unexpected given the quality of the *B. mori* genome assembly and the potential age of this duplication event. Thus, these results are consistent with some type of ancient duplication event in the ancestry of the Lepidoptera.

### Biased Gene Retention and Loss Following Inferred WGDs

Given the rarity of polyploidy among insects and the fact that no ancient WGDs have been previously observed in the clade, we further characterized the nature of these putative WGDs. A common signature of paleopolyploidy is the biased retention and loss of genes relative to the background pattern of gene turnover. Surviving paralogs from ancient WGDs are often enriched with idiosyncratic gene ontology (GO) categories in plants (28, 37–40), yeasts (41), and animals (1, 42, 43). Among the many hypotheses to explain the retention of paralogs (44–46), the dosage balance hypothesis (DBH) is the only hypothesis that predicts the parallel retention and loss of functionally related genes following WGDs across species (45, 47, 48). The DBH predicts that genes with many connections or in dosage sensitive regulatory networks will be retained in duplicate following polyploidy to maintain the relative abundance of protein products. Conversely, these same genes will be preferentially lost following small-scale gene duplications to prevent the disruption of dosage (45). Thus, if the signatures of gene duplication observed in the insects are the result of WGDs, we expect to find a biased pattern of retention and loss that may be shared among the putative insect WGDs.

To test for biased gene retention and loss among our inferred WGDs, we used a HMMR based approach to annotate the genomes or transcriptomes of 20 hexapod and one outgroup species with the *D. melanogaster* Gene Ontology (GO) data (49). Paralogs were partitioned from the putative WGDs inferred in Ks plots by fitting a normal mixture model to the distributions (SI Appendix, Table S5). Using the numbers of genes annotated to each GO category, we performed a principal component analysis to assess the overall differences in GO category composition among all genes in the genome/transcriptome and paralogs retained from each putative WGD (Fig. 3, SI Appendix, Dataset S7). These categories of genes formed non-overlapping clusters in the PCA. Notably, the GO composition of WGD paralogs from each species formed a narrower confidence interval than the entire transcriptomes/genomes. Significant differences (*p* < 0.001) between the GO composition of these two groups were also found using a chi-square goodness of fit test. Paralogs from WGDs across all 21 species demonstrated biased patterns of gene retention and loss (Fig. 3, SI Appendix, Fig. S13-S14). Consistent with our PCA, many of the same GO categories were significantly over- and under-represented among the genes maintained in duplicate from the putative insect WGDs. For example, over 50% of our sampled species had paralogs from the ancient WGDs that were significantly enriched for RNA metabolic processes, nucleus component, DNA binding function, and nucleobase-containing compound metabolic process(SI Appendix, Fig. S13). Similarly, many GO categories were significantly under-retained among the paralogs of ancient WGDs (SI Appendix, Fig. S13). Over half of our sampled species demonstrated significant under-retention of genes associated with oxidoreductase activity, hydrolase activity, electron carrier activity, peptidase activity, and proteolysis. Although there is some noise in the patterns of gene retention, this may not be surprising given the great divergence of many species from the annotation source, *Drosophila*, and the phylogenetic scale of the hexapods. Considering the great diversity of these lineages, the convergent pattern of gene retention and loss is consistent with our expectations following polyploidy.

**Fig. 3.**
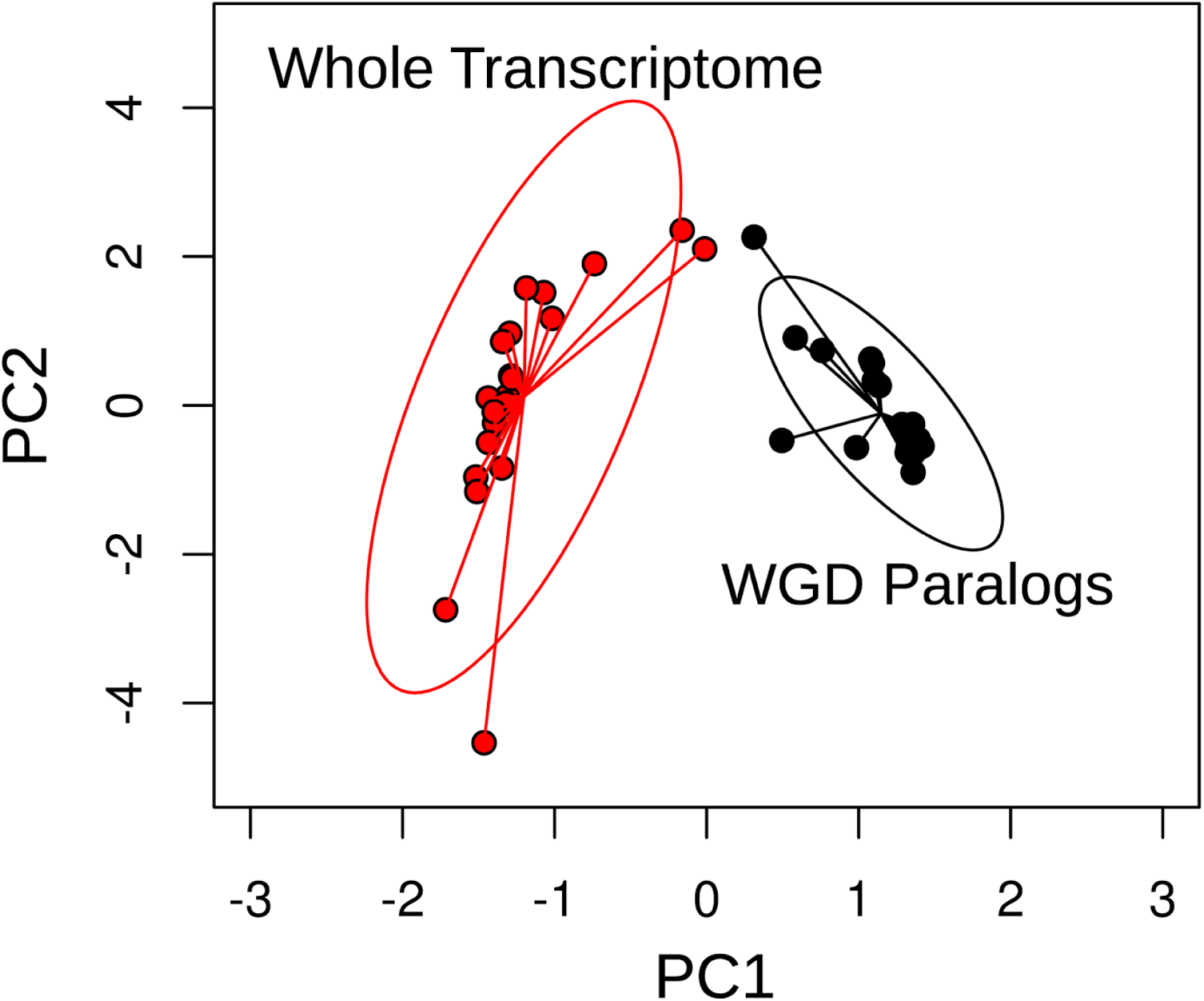
Principal component analysis of the GO category composition of all genes in each genome/transcriptome and WGD paralogs. Red circles = number of genes annotated to each GO category in the whole genome or transcriptomes. Black circles = number of WGD paralogs annotated to each GO category. Ellipses represent the 95% confidence interval of standard deviation of point scores.

## Discussion

Our analyses provide the first evidence for paleopolyploidy in the hexapods. Combining our gene age distribution and phylogenomic analyses, we found evidence for 24 significant, episodic bursts of gene duplication in the insects. Although some of these duplication events may result from other mechanisms of gene duplication, they appear to be consistent with WGDs inferred using similar approaches in plants (28, 50, 51) and animals (1, 24). Of these bursts, 18 were detected as peaks in gene age distributions that are characteristic of ancient WGDs in 14 hexapod orders (Fig. 2, SI Appendix, Table S3). Genes retained in duplicate from these 18 putative WGD events had a shared pattern of biased gene retention and loss, an expected result of paleopolyploidy but not other types of gene duplication (44, 45). An additional six large-scale genome duplication events were inferred deeper in the phylogeny of insects using phylogenomic analyses. These six duplications were consistent with simulated WGDs at these locations in the insect phylogeny. Many phylogenomic analyses use a single value of gene duplication number per branch to diagnose a WGD (52, 53). However, variation in branch lengths and gene birth/death rates may confound these phylogenomic inferences of gene duplication (54). The number of duplications on a given branch is expected to covary with branch length. Without taking into account branch length variation, the number of duplications may appear to change dramatically from branch to branch. Our use of simulated gene trees with both null and positive simulations of WGDs should provide a more robust inference of large-scale genome duplication events that is less sensitive to branch length variation than previous approaches (8, 52, 53). Although many of these inferred duplications are consistent with simulated WGDs, we refer to them here as large-scale duplication events given the phylogenetic depth and difficulty assessing their nature with other methods. Notably, we do find statistical evidence from analyses of synteny in the *B. mori* genome that a duplication in the ancestry of the Lepidoptera likely involved structural duplication rather than just gene duplication alone. However, more complete genomic analyses are needed to confirm the nature of these duplication events.

Our analyses of GO categories revealed a largely convergent pattern of biased gene retention and loss following genome duplication in multiple species of insects. Based on the dosage balance hypothesis (DBH), we expected to observe biased retention and loss of similar GO categories following WGDs rather than other types of gene duplication events (44, 45). Our observation across multiple species is consistent with post-polyploid genome evolution because dosage-balance may drive the convergent retention of genes. Although neo- and subfunctionalization are also important in duplicate gene retention, only the DBH predicts a convergent pattern of gene retention following polyploidy across multiple species (44, 45). Thus, the signal of convergent gene retention is consistent with our inference that they were likely WGDs. This biased pattern was also found among WGD paralogs in an outgroup species, *Ixodes scapularis* (deer tick), and suggests that it may be consistent across arthropods. Previous studies in plants have found that similar functional categories of genes have been maintained following different genome duplication events (38, 40), although there are often idiosyncratic patterns observed across families (28, 37, 38). The 20 insect transcriptomes included in our GO category analyses represent diverse hexapod (SI Appendix, Fig. S13) orders whose divergence times far exceed most previously studied plant examples. At least among the arthropods, our analyses suggest that biases in duplicate gene retention may be maintained over hundreds of millions of years. A potential explanation for the long consistency of this pattern is that insects have experienced a limited number of ancient WGDs that may influence large-scale shifts in gene network relationships. Most hexapod species included in our analyses only had one round of genome duplication, whereas nearly all flowering plants have likely experienced at least three rounds of paleopolyploidy(2).

Our discovery of WGDs and other large-scale genome duplications in the ancestry of hexapods raises many questions about the role of gene and genome duplication in plant and animal evolution. It has long been known that polyploidy is rarer in animals than in plants (16, 17). Mueller hypothesized that sex chromosomes are barriers to polyploidy in animals (16, 55). Although our sample size is limited, we observed some patterns in our data set consistent with this hypothesis. For example, we observed more putative WGDs in the Trichoptera, with Z0 sex-determination, relative to its sister lineage, the Lepidoptera, which mostly has a ZW system (56, 57). Although we had a small sample size for each lineage, this observation is expected because WGD will cause less disruption of dosage compensation in Z0 compared to ZW systems (16, 17).

More refined placement and denser sampling of ancient WGDs in insects will provide a new opportunity to test Mueller’s classic hypothesis. Similarly, a recent study proposed that the phylogenetic distribution of ancient WGD among plants may be a by-product of asexuality rather than an intrinsic advantage of polyploidy itself (58). Given the diversity of insect sexual systems, a better understanding of ancient WGDs in insects would provide an improved context for testing this hypothesis in other eukaryotes. Our results also lend support to long-standing hypotheses that gene and genome duplications are important forces in animal evolution (19, 59) in insects such as the expansion of *Hox* genes in the Lepidoptera (60) and during the co-evolutionary radiation of pierid butterflies and the Brassicales (10). Our observation of numerous episodes of duplication in the hexapod phylogeny raises the possibility that large-scale duplications may be associated with the evolution of novelty and diversity across the insect phylogeny, including additional duplication driven co-evolutionary interactions with plants. Further phylogenomic sampling of insects will likely reveal more paleopolyploidy and other large-scale genome duplications. Large sequencing projects such as 1KP (61), 1KITE (26), and i5K (62) will improve our ability to place these events in the plant and hexapod phylogenies. As it stands, our results indicate that large-scale gene and genome duplications have occurred during the evolution of the most diverse clade of eukaryotes.

## Materials and Methods

### Data sampling

We compiled a phylogenetically diverse genomic dataset that comprised every hexapod order and outgroups from related arthropods (SI Appendix, Dataset S1). These data included 119 transcriptomes and 25 genomes for hexapods, as well as nine transcriptomes and two genomes from the Chelicerates, myriapods, and crustaceans as outgroups. We downloaded 128 published transcriptome assemblies from the GenBank Transcriptome Shotgun Assembly database (TSA), and 27 published genomes from multiple genome databases (SI Appendix).

### DupPipe: inference of WGDs from paralog age distributions

For each data set, we used our DupPipe pipeline to construct gene families and estimate the age of gene duplications (27). We translated DNA sequences and identified reading frames by comparing the Genewise alignment to the best hit protein from a collection of proteins from 24 metazoan genomes from Metazome v3.0. For each node in our gene family phylogenies, we estimated synonymous divergence (Ks) using PAML with the F3X4 model (63). For each species, we identified ancient WGDs as significant peaks of gene duplication in histograms of the age distribution of gene duplications (Ks plots) using mixture models (64) and SiZer (65).

### MAPS: phylogenomic inference of large-scale genome duplications from nuclear gene trees

To infer large-scale genome duplications, we used the Multi-tAxon Paleopolyploidy Search (MAPS) tool (8), a gene tree sorting and counting algorithm. We translated each transcriptome into amino acid sequences using the TransPipe pipeline (27). Using these translations, we performed reciprocal BLASTP searches for each MAPS data set with an E-value of 10e-5 as a cutoff. We clustered gene families from these BLAST results using OrthoMCL v2.0 with the default parameters (66) and only retained gene families that contained at least one gene copy from each taxon. We used SATé for alignment and phylogeny reconstruction of gene families (67). The best scoring SATé tree for each gene family was used to infer large-scale genome duplications with MAPS. Results of all 33 MAPS analyses are provided in Dataset S3.

### *Syntenic analysis of* Bombyx mori

To validate our MAPS inferences of large-scale genome duplications in the Lepidoptera, we examined the the *Bombyx mori* genome (31) for syntenic evidence of duplication (SI Appendix, Figs. S6*AA*-S9*AA*). SynMap on the CoGe platform (35) was used to identify syntenic regions. We used blastp with E-value cutoff of 10e-5. Quota Align Merge was selected to merge syntenic chains and other parameters were set as default. Synteny was detected with a minimum of three genes to seed a chain and a Manhattan distance of 40.

### Estimating orthologous divergence to place large-scale genome duplications in relation to lineage divergence

We estimated ortholog divergences among major hexapod clades to place large-scale genome duplications in relation to lineage divergence (SI Appendix, Table S2). We used the RBH Ortholog pipeline (27) to estimate the median ortholog divergence between 24 species pairs (SI Appendix, Fig. S5 and Table S2). The median ortholog divergence was used to estimate the lower bound of paralog divergence for shared ancient large-scale genome duplication events.

### Gene Ontology (GO) annotations and paleolog retention and loss patterns

We used the best hit with length of at least 100 bp and an E-value of at least 0.01 in phmmer (HMMER 3.1b1) for Gene Ontology (GO) annotation. For each species, we assigned paralogs to ancient WGDs based on the Ks ranges identified in mixture model analyses (SI Appendix, Table S5) (28, 64). We evaluated the overall differences between the genome/transcriptomes and WGD paralogs by performing a principal component analysis (PCA) using the rda function in R package vegan (68). We also tested for differences among GO annotations across the inferred WGD events using chi-square tests (SI Appendix, Fig. S13-S14).

## Acknowledgments

We thank A. Baltzell, A. Baniaga, J. Bronstein, K. Dlugosch, S. Jorgensen, E. Lyons, X. Qi, J. Czekanski-Moir and Y. Tomoyasu for suggestions and discussion. Z. Wang, L. Tang, and X. Song for assistance with insect images. We also thank two anonymous reviewers for their careful reading and suggestions that improved the manuscript. Hosting infrastructure and services provided by the Biotechnology Computing Facility (BCF) at the University of Arizona. M.S.B. was supported by NSF-IOS-1339156 and NSF-EF-1550838.

## References

1. Berthelot C, et al. (2014) The rainbow trout genome provides novel insights into evolution after whole-genome duplication in vertebrates. Nat Commun 5:3657.

2. Barker MS, Husband BC, Pires JC (2016) Spreading Winge and flying high: The evolutionary importance of polyploidy after a century of study. Am J Bot 103(7):1139–1145.

3. Van de Peer Y, Mizrachi E, Marchal K (2017) The evolutionary significance of polyploidy. Nat Rev Genet. 18(7):411–424.

4. Wolfe KH, Shields DC (1997) Molecular evidence for an ancient duplication of the entire yeast genome. Nature 387(6634):708–713.

5. Wood TE, et al. (2009) The frequency of polyploid speciation in vascular plants. Proc Natl Acad Sci USA 106(33):13875–13879.

6. Barker MS, Arrigo N, Baniaga AE, Li Z, Levin DA (2016) On the relative abundance of autopolyploids and allopolyploids. New Phytol 210(2):391–398.

7. Jiao Y, et al. (2011) Ancestral polyploidy in seed plants and angiosperms. Nature 473(7345):97–100.

8. Li Z, et al. (2015) Early genome duplications in conifers and other seed plants. Science Advances 1(10):e1501084.

9. Wendel JF (2015) The wondrous cycles of polyploidy in plants. Am J Bot 102(11):1753–1756.

10. Edger PP, et al. (2015) The butterfly plant arms-race escalated by gene and genome duplications. Proc Natl Acad Sci USA 112(27):8362–8366.

11. Gout J-F, Lynch M (2015) Maintenance and loss of duplicated genes by dosage subfunctionalization. Mol Biol Evol. 32(8):2141–2148.

12. Arrigo N, Barker MS (2012) Rarely successful polyploids and their legacy in plant 1. genomes. Curr Opin Plant Biol 15(2):140–146.

13. Murat F, Armero A, Pont C, Klopp C, Salse J (2017) Reconstructing the genome of the most recent common ancestor of flowering plants. Nat Genet. 49(4):490–496.

14. Selmecki AM, et al. (2015) Polyploidy can drive rapid adaptation in yeast. Nature. 519(7543):349–352.

15. Tank DC, et al. (2015) Nested radiations and the pulse of angiosperm diversification: increased diversification rates often follow whole genome duplications. New Phytol 207(2):454–467.

16. Orr HA (1990) “Why Polyploidy is Rarer in Animals Than in Plants” Revisited. Am Nat 136(6):759–770.

17. Otto SP, Whitton J (2000) Polyploid incidence and evolution. Annu Rev Genet 34:401–437.

18. McLysaght A, Hokamp K, Wolfe KH (2002) Extensive genomic duplication during early chordate evolution. Nat Genet 31(2):200–204.

19. Ohno S (1970) Evolution by Gene Duplication (Springer, New York).

20. Gregory TR, Mable BK (2005) Polyploidy in animals. The Evolution of the Genome, eds Gregory TR (Elsevier Academic, London), pp 427–517.

21. Hallinan NM, Lindberg DR (2011) Comparative analysis of chromosome counts infers three paleopolyploidies in the Mollusca. Genome Biol Evol 3:1150–1163.

22. Nossa CW, et al. (2014) Joint assembly and genetic mapping of the Atlantic horseshoe crab genome reveals ancient whole genome duplication. Giga Sci 3(1):9.

23. Clarke TH, Garb JE, Hayashi CY, Arensburger P, Ayoub NA (2015) Spider transcriptomes identify ancient large-scale gene duplication event potentially important in silk gland evolution. Genome Biol Evol. 7(7):1856–1870.

24. Schwager EE, et al. (2017) The house spider genome reveals an ancient whole-genome duplication during arachnid evolution. BMC Biol 15(1):62.

25. De Wever A. eds. (2015) Species 2000 & ITIS Catalogue of Life, 2015 Annual Checklist. Catalogue of Life. Available at: www.catalogueoflife.org/annual-checklist/2015 [Accessed October 20, 2015].

26. Misof B, et al. (2014) Phylogenomics resolves the timing and pattern of insect evolution. Science 346(6210):763–767.

27. Barker MS, et al. (2010) EvoPipes.net: Bioinformatic Tools for Ecological and Evolutionary Genomics. Evol Bioinform Online 6:143–149.

28. Barker MS, et al. (2008) Multiple paleopolyploidizations during the evolution of the Compositae reveal parallel patterns of duplicate gene retention after millions of years. Mol Biol Evol 25(11):2445–2455.

29. SjÖstrand J, Arvestad L, Lagergren J, Sennblad B (2013) GenPhyloData: realistic simulation of gene family evolution. BMC Bioinformatics 14:209.

30. Rabier C-E, Ta T, Ané C (2014) Detecting and locating whole genome duplications on a phylogeny: a probabilistic approach. Mol Biol Evol 31(3):750–762.

31. International Silkworm Genome Consortium (2008) The genome of a lepidopteran model insect, the silkworm *Bombyx mori*. Insect Biochem Mol Biol 38(12):1036–1045.

32. Cutter AD (2008) Divergence times in *Caenorhabditis* and *Drosophila* inferred from direct estimates of the neutral mutation rate. Mol Biol Evol 25(4):778–786.

33. Vanneste K, Van de Peer Y, Maere S (2013) Inference of genome duplications from age distributions revisited. Mol Biol Evol 30(1):177–190.

34. Cui L, et al. (2006) Widespread genome duplications throughout the history of flowering plants. Genome Res 16(6):738–749.

35. Lyons E, Pedersen B, Kane J, Freeling M (2008) The value of nonmodel genomes and an example using SynMap within CoGe to dissect the hexaploidy that predates the rosids. Trop Plant Biol 1(3-4):181–190.

36. Nakazato T, Jung M-K, Housworth EA, Rieseberg LH, Gastony GJ (2006) Genetic map-based analysis of genome structure in the homosporous fern *Ceratopteris richardii*. Genetics 173(3):1585–1597.

37. Mandáková T, Li Z, Barker MS, Lysak MA (2017) Diverse genome organization following 13 independent mesopolyploid events in Brassicaceae contrasts with 1. convergent patterns of gene retention. Plant J. 91(1):3–21.

38. Li Z, et al. (2016) Gene duplicability of core genes Is highly consistent across all angiosperms. Plant Cell 28(2):326–344.

39. De Smet R, et al. (2013) Convergent gene loss following gene and genome duplications creates single-copy families in flowering plants. Proc Natl Acad Sci USA 110(8):2898–2903.

40. Rody HVS, Baute GJ, Rieseberg LH, Oliveira LO (2017) Both mechanism and age of duplications contribute to biased gene retention patterns in plants. BMC Genomics 18(1):46.

41. Conant GC (2014) Comparative genomics as a time machine: how relative gene dosage and metabolic requirements shaped the time-dependent resolution of yeast polyploidy. Mol Biol Evol 31(12):3184–3193.

42. Session AM, et al. (2016) Genome evolution in the allotetraploid frog *Xenopus laevis*. Nature 538(7625):336–343.

43. Lien S, et al. (2016) The Atlantic salmon genome provides insights into rediploidization. Nature 533(7602):200–205.

44. Kondrashov FA, Kondrashov AS (2006) Role of selection in fixation of gene duplications. J Theor Biol 239(2):141–151.

45. Freeling M (2009) Bias in plant gene content following diffierent sorts of duplication: tandem, whole-genome, segmental, or by transposition. Annu Rev Plant Biol 60(1):433–453.

46. Hahn MW (2009) Distinguishing among evolutionary models for the maintenance of gene duplicates. J Hered 100(5):605–617.

47. Freeling M, Thomas BC (2006) Gene-balanced duplications, like tetraploidy, provide predictable drive to increase morphological complexity. Genome Res 16(7):805–814.

48. Conant GC, Birchler JA, Pires JC (2014) Dosage, duplication, and diploidization: clarifying the interplay of multiple models for duplicate gene evolution over time. Curr Opin Plant Biol 19:91–98.

49. Gene Ontology Consortium (2015) Gene Ontology Consortium: going forward. 1. Nucleic Acids Res 43(Database issue):D1049–1056.

50. Barker MS, et al. (2016) Most Compositae (Asteraceae) are descendants of a paleohexaploid and all share a paleotetraploid ancestor with the Calyceraceae. Am J Bot 103(7):1203–1211.

51. Badouin H, et al. (2017) The sunflower genome provides insights into oil metabolism, flowering and Asterid evolution. Nature 546(7656):148–152.

52. Huang C-H, et al. (2016) Multiple polyploidization events across Asteraceae with two nested events in the early history revealed by nuclear phylogenomics. Mol Biol Evol 33(11):2820–2835.

53. Yang Y, et al. (2015) Dissecting molecular evolution in the highly diverse plant clade Caryophyllales using transcriptome sequencing. Mol Biol Evol 32(8):2001–2014.

54. Hahn MW (2007) Bias in phylogenetic tree reconciliation methods: implications for vertebrate genome evolution. Genome Biol 8(7):R141.

55. Muller HJ (1925) Why polyploidy is rarer in animals than in plants. Am Nat 59(663):346–353.

56. Blackmon H, Ross L, Bachtrog D (2017) Sex determination, sex chromosomes, and karyotype evolution in Insects. J Hered 108(1):78–93.

57. Tree of Sex Consortium (2014) Tree of Sex: a database of sexual systems. Sci Data 1:140015.

58. Freeling M (2017) Picking up the ball at the K/Pg boundary: the distribution of ancient polyploidies in the plant phylogenetic tree as a spandrel of asexuality with occasional sex. Plant Cell 29(2):202–206.

59. Van de Peer Y, Maere S, Meyer A (2009) The evolutionary significance of ancient genome duplications. Nat Rev Genet 10(10):725–732.

60. Ferguson L, et al. (2014) Ancient expansion of the hox cluster in lepidoptera generated four homeobox genes implicated in extra-embryonic tissue formation. PLoS Genet 10(10):e1004698.

61. Wickett NJ, et al. (2014) Phylotranscriptomic analysis of the origin and early 1. diversification of land plants. Proc Natl Acad Sci USA 111(45):E4859–E4868.

62. K Consortium (2013) The i5K Initiative: advancing arthropod genomics for knowledge, human health, agriculture, and the environment. J Hered 104(5):595–600.

63. Yang Z (1997) PAML: a program package for phylogenetic analysis by maximum likelihood. Bioinformatics 13(5):555–556.

64. McLachlan GJ, Peel D (1999) The EMMIX algorithm for the fitting of normal andt-Components. J Stat Softw 4(2):1–14.

65. Chaudhuri P, Marron JS (1999) SiZer for exploration of structures in curves. J Am Stat Assoc 94(447):807–823.

66. Li L, Stoeckert CJ Jr, Roos DS (2003) OrthoMCL: identification of ortholog groups for eukaryotic genomes. Genome Res 13(9):2178–2189.

67. Liu K, Raghavan S, Nelesen S, Linder CR, Warnow T (2009) Rapid and accurate large-scale coestimation of sequence alignments and phylogenetic trees. Science 324(5934):1561–1564.

68. Dixon P (2003) VEGAN, a package of R functions for community ecology. J Veg Sci 14(6):927.

